# Single-cell transcriptomics reveals subset-specific metabolic profiles underpinning the bronchial epithelial response to flagellin

**DOI:** 10.1101/2022.11.02.514918

**Authors:** Ivan Ramirez-Moral, Alex R. Schuurman, Joe M. Butler, Xiao Yu, Karen de Haan, Sarah van Leeuwen, Alex F. de Vos, Menno D. de Jong, Felipe A. Vieira Braga, Tom van der Poll

**Author notes:** These authors contributed equally to this work (Ivan Ramirez-Moral, Alex R. Schuurman). These authors also contributed equally to this work (Felipe A. Vieira Braga, Tom van der Poll). **Corresponding author**: Ivan Ramirez-Moral, Amsterdam UMC, location Academic Medical Center, University of Amsterdam, Center of Experimental and Molecular Medicine, Meibergdreef 9, Room T10-236, 1105 AZ Amsterdam, The Netherlands.

## Abstract

Respiratory epithelial cells line the airways and represent the first line of defense against respiratory pathogens. The cellular heterogeneity of the airway wall has only recently been recognized by single-cell analyses. Here, we leveraged single-cell RNA sequencing of primary human bronchial epithelial cells growing in air-liquid interface to determine cell-specific changes evoked by flagellin, a protein driving the motility of many mucosal pathogens. We detected seven cell clusters in the human epithelium, including ciliated cells, ionocytes and several states of basal and secretory cells, of which only inflammatory basal cells and inflammatory secretory cells showed a proportional increase in response to flagellin. Only inflammatory secretory cells showed evidence of metabolic reprogramming toward aerobic glycolysis, and inhibition of the mTOR pathway specifically reduced this subset, prevented this metabolic shift, and reduced inflammatory gene transcription in these cells. This study expands our knowledge of the airway’s immune response to flagellated pathogens to single cell resolution and defines a novel target to modulate mucosal immunity during bacterial infections.

## INTRODUCTION

The human airways represent a major interface between the environment, which is teeming with potentially pathogenic microbes, and the inner body. The epithelium that lines the airways is responsible for maintaining immune homeostasis in the lung, while also offering protection to microorganisms and pollutants that we inhale^1,2^. Over the last decade, the field of immunometabolism has solidified the concept that regulation of the different energy pathways within cells is key in maintaining and shaping the immune response to infection^3–6^. We recently engaged in studies on the role of metabolism in the immune function of the airway’s epithelium^7,8^, and showed that activation of primary human bronchial epithelial (HBE) cells in response to flagellin – a bacterial component that is typically present on the surface of mucosal pathogens^9,10^ – is regulated by mammalian target of rapamycin (mTOR)-driven glycolysis^7^. Knowledge of the mechanisms by which flagellin activates the respiratory epithelium is not only of relevance for understanding local inflammatory responses induced by flagellated pathogens, but also considering that administration of purified flagellin via the airways has been suggested as an immune enhancing therapy during infection of the lungs^9,11^. The human airway epithelium consists of a heterogeneous community of different cell types, each with unique molecular characteristics^2^. While it is likely that these different cell types, which fulfill distinct functions during the immune response against infection, are driven by specific metabolic programs, this remains to be investigated.

In this study we sought to build upon our previous findings by analyzing the epithelial response to flagellin at a higher resolution. Single-cell RNA sequencing (scRNAseq) makes it possible to individually analyze thousands of cells from a complex system, such as the airway epithelium, and to identify cell subsets with unique transcriptomic profiles. We applied this approach to our airway model of primary HBE cells growing at the air-liquid interface. Through scRNAseq we identified seven different cell clusters – including ionocytes, ciliated cells, and several states of basal and secretory cells – and observed that inflammatory subsets of basal and secretory cells were more abundant in HBE cell cultures exposed to flagellin. Interestingly, we confirmed flagellin-induced metabolic reprogramming towards aerobic glycolysis, but this was only present in secretory inflammatory cells. Inhibition of the mTOR pathway with rapamycin prevented this flagellin-induced metabolic shift, resulted in strongly decreased proportions of secretory inflammatory cells, and suppressed the inflammatory transcriptomic response of secretory cells. These data expand upon our previous findings and reveal how specific epithelial cell subsets rely on different metabolic pathways in their response to flagellin. Specifically, inhibition of the mTOR pathway appears to present a specific tool for limiting the inflammatory response of secretory epithelial cells, which may offer a targeted therapeutic approach for controlling inflammation in the lungs.

## RESULTS

HBE cells were harvested from healthy tracheobronchial tissue obtained during lobectomy surgery and differentiated in the air-liquid interface as previously described^7^. This approach yielded fully differentiated HBE cells constituting three cell types – basal, secretory, and ciliated cells – in a basal-to-apical orientation as shown by confocal microscopy (**Figure 1A**). After quality-control we analyzed the transcriptome of 6577 single HBE cells across three experimental conditions: unstimulated (vehicle; PBS), stimulated with flagellin, or flagellin-stimulated in the presence of rapamycin.

**Figure. 1.**
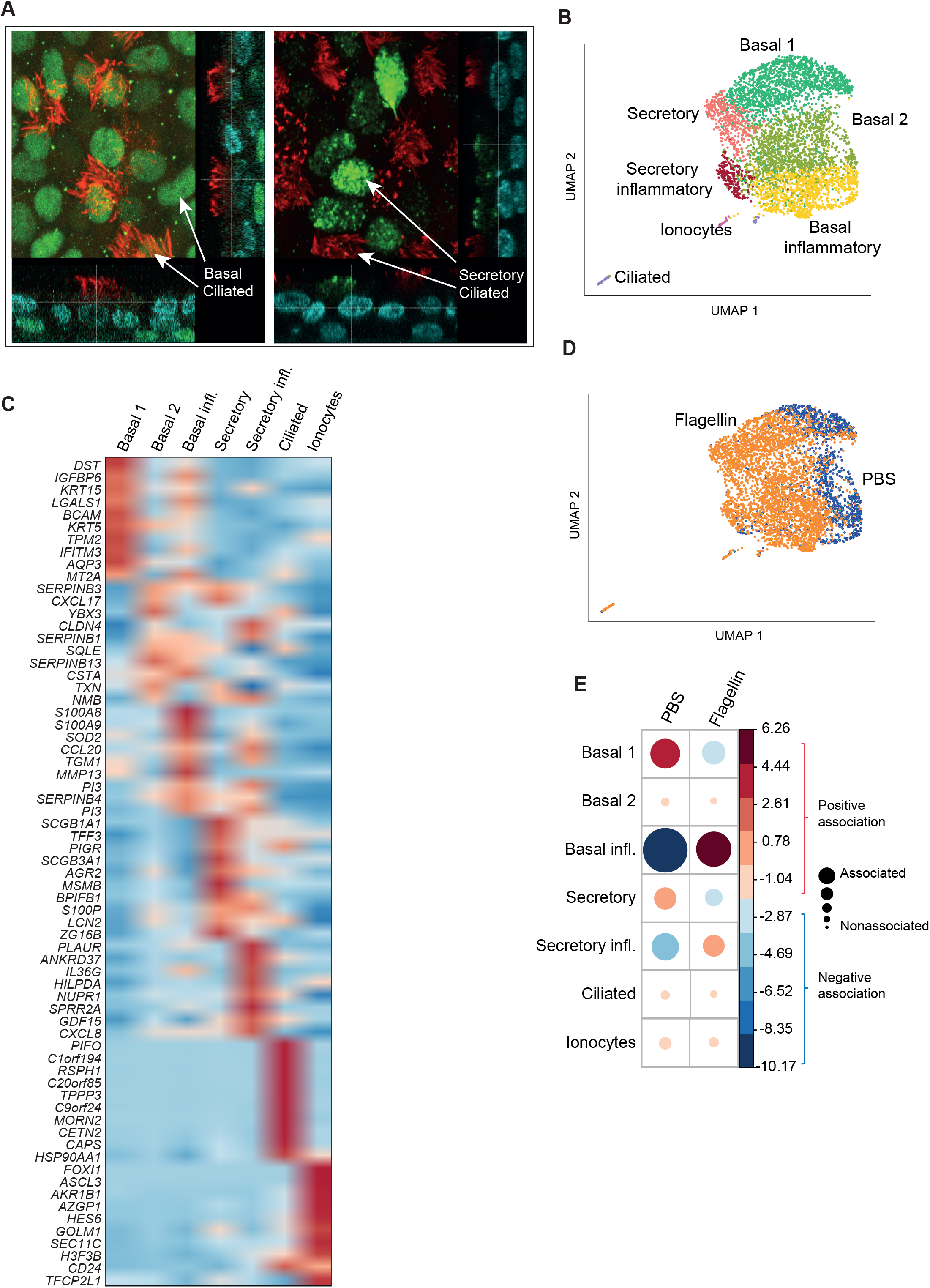
Primary human bronchial epithelial cell cultures show a proportional increase of basal inflammatory and secretory inflammatory cells upon stimulation with flagellin. **A**) Confocal imaging of polarized HBE cells; in red β-tubulin^+^ ciliated cells, blue nuclei (DAPI), green (left panel) p63^+^ basal cells and green (right panel) MUC5B^+^ secretory cells. Max. projection. **B**) UMAP of HBE cells, stimulated by either PBS or flagellin, where the color indicates the cluster as identified by single-cell transcriptomic analysis. Each dot represents one cell. Genes from the canonical markers used for cluster identification are listed in **Table S1. C**) Heatmap showing the expression of the top differentially expressed genes (DEGs) for each cluster, when compared to the rest of the clusters. **D**) Same as panel B, now colored by experimental condition. **E**) Correlation plot depicting cluster enrichment between the PBS and flagellin condition. Dot size is proportional to the Pearson’s residual of the chi-squared test (reflecting the difference between the observed and expected proportion), while the color represents the degree of association from Pearson’s chi-squared residuals (red is a positive association, blue is a negative association)

### Increased proportions of inflammatory basal and secretory cells upon stimulation with flagellin

We first explored the epithelial response to flagellin when compared to unstimulated cells. Based on the transcriptome of all individual cells from these two conditions we identified seven cell clusters, as visualized by Uniform Manifold Approximation and Projection (UMAP) dimensionality reduction (**Figure 1B**). Clusters were annotated based on canonical marker genes among the top differentially expressed genes (DEGs) between clusters (see **Supplemental Table 1** for an overview^2,12^), which revealed three clusters of basal cells and two clusters of secretory (goblet) cells, as well as ionocytes and ciliated cells (**Figure 1B**). Two of the three basal cell states reflected different differentiation stages: relatively high *KRT5* and *KRT15* expression suggests that basal 1 cells were more stem-like, while basal 2 cells were more mature. The third basal subset we labelled as inflammatory based on high expression of S100 members *S100A8* and *S100A9* and other inflammatory mediators such as *CCL20*. Secretory cells were characterized by high expression of the canonical markers^2,12^ *SCGB1A1, MUC5B* and *SCGB3A1*. An inflammatory secretory subset was identified based on high expression of the genes encoding for the inflammatory mediators CXCL8, CXCL16 and G-CSF, among others. Ciliated cells were identified by high levels of the canonical marker genes *PIFO* and *FOXJ1*. Ionocytes were defined by high expression of *FOXI1*. The top 10 DEGs, not only including canonical genes, between the identified clusters are depicted in **Figure 1C**. Next, we assessed the effect of flagellin on the composition of the HBE cells (**Figure 1D** shows the UMAP colored per condition). Flagellin induced profound differences in the proportional composition of the identified cell clusters. Specifically, we found a lower proportion of basal 1 and secretory cells, and a higher proportion of basal inflammatory and secretory inflammatory cells in the flagellin condition (**Figure 1E**). We observed no differences in the proportion of basal 2 cells, ciliated cells, or ionocytes after flagellin stimulation. As the ciliated cell cluster and ionocyte cluster represented very few cells and showed no proportional change in the flagellin condition, we focused on the basal and secretory cell clusters.

### The transcriptional landscape of inflammatory basal and secretory cells

As only the inflammatory subclusters of the basal and secretory cells were clearly more abundant in the flagellin condition, we next investigated the transcriptional profile of these inflammatory subclusters. We compared the secretory inflammatory cells to the other secretory cells, and showed that genes coding for the prototypic epithelial-inflammatory mediators were up-regulated in the secretory inflammatory cells (*CCL20, CXCL8, CSF3*) as well as three keratin coding genes (*KRT17, KRT15, KRT16*) (**Figure 2A**). Interestingly, the gene coding for the enzyme responsible of the last reaction of the glycolysis pathway (*LDHA*) was highly expressed in the secretory inflammatory subset (**Figure 2A**). Basal inflammatory cells, relative to basal 1 cells, displayed a strong pro-inflammatory signature indicative of a heightened innate immune response, including upregulation of *S100A8-9*, several *SERPIN* genes, *CXCL1, CXL8* and *CCL20* (**Figure 2B**, see **Supplemental Figure 1** for basal inflammatory versus basal 2). Downregulated genes included two keratin coding genes (*KRT5, KRT15*), and two tropomyosin coding genes (*TPM1, TPM2*). Next, we sought to identify unique features between these two inflammatory subclusters. The Venn-Euler plot (**Figure 2C**) shows the result from the differential gene analysis directly comparing these subclusters, indicating the number of genes significantly upregulated in secretory inflammatory cells (255), upregulated in basal inflammatory cells (565) and non-significant genes (233). We performed pathway analysis on the 255 upregulated genes in secretory inflammatory cells, and the 565 upregulated genes in basal inflammatory cells (**Figure 2C**). Several of the upregulated pathways – such as hypoxia, the P53 pathway and TNF signaling via NFkB – were enriched in both cell types. Interestingly, only the secretory inflammatory cells had upregulated glycolysis and MTOR signaling pathways, while basal inflammatory cells showed upregulated transcripts involved in oxidative phosphorylation. While this was in line with our previous findings, in which we reported upregulated mTOR-mediated glycolysis in bulk HBE cells stimulated by flagellin^7^, the present data suggested that this metabolic switch may specifically relate to inflammatory secretory cells. To further investigate this, we next analyzed the metabolic programming of the identified clusters, and the effect of flagellin thereon.

**Figure 2.**
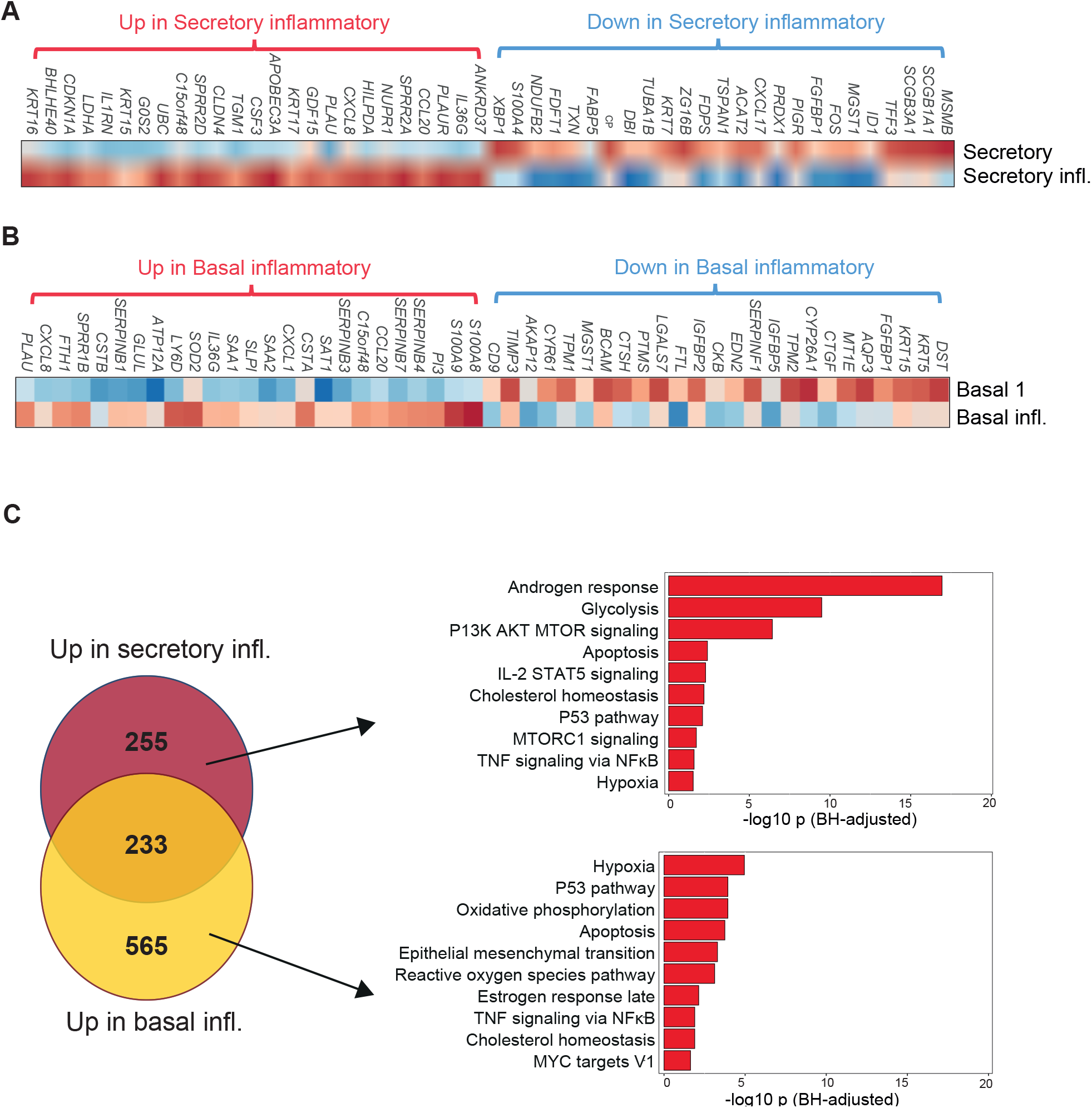
The transcriptional landscape of inflammatory basal and secretory cells. **A**) Heatmap showing the expression of the top 25 increased and top 25 decreased DEGs derived from comparing the secretory and secretory inflammatory cells (log fold change > 0.25, adjusted p<0.05). **B**) Same as in panel A, but here the comparison is made between basal 1 and basal inflammatory cells. **C**) Venn-Euler plot depicting the result of differential gene expression analysis between secretory inflammatory and basal inflammatory cells. Molecular Signatures Database (MSigDB) pathway analysis was performed on the DEGs upregulated in secretory inflammatory cells (upper pathways), and on the DEGs upregulated in basal inflammatory cells (lower pathways). The X-axis shows the Benjamini-Hochberg adjusted −log10 p-value from the enrichment score analysis.

### Secretory inflammatory cells have a distinct metabolic profile of enhanced glycolysis and mTOR-signaling

We compared metabolic signature scores for amino acid metabolism, glycolysis, mTOR signaling, lipid metabolism and the tricarboxylic acid (TCA) cycle – derived from the hallmark gene sets in the Molecular Signatures Database^13^ – between all basal and secretory clusters. In line with the analysis in **Figure 2C**, we found glycolysis and mTOR signaling to be clearly upregulated in secretory inflammatory cells when compared to the other subsets, including the other secretory cells (**Figure 3A**). The other metabolic pathways showed no major differences between subsets, although genes encoding for enzymes of the TCA cycle were relatively lowly expressed in secretory inflammatory cells and highly expressed in the other secretory subset, possibly indicating a metabolic shift from the TCA cycle to glycolysis, triggered by flagellin (**Supplemental Figure 2**). Heatmaps of key genes in glycolysis and mTOR-signaling corroborated the gene signature scores, with clearly increased expression specifically in secretory inflammatory cells (**Figure 3B**).

**Figure 3.**
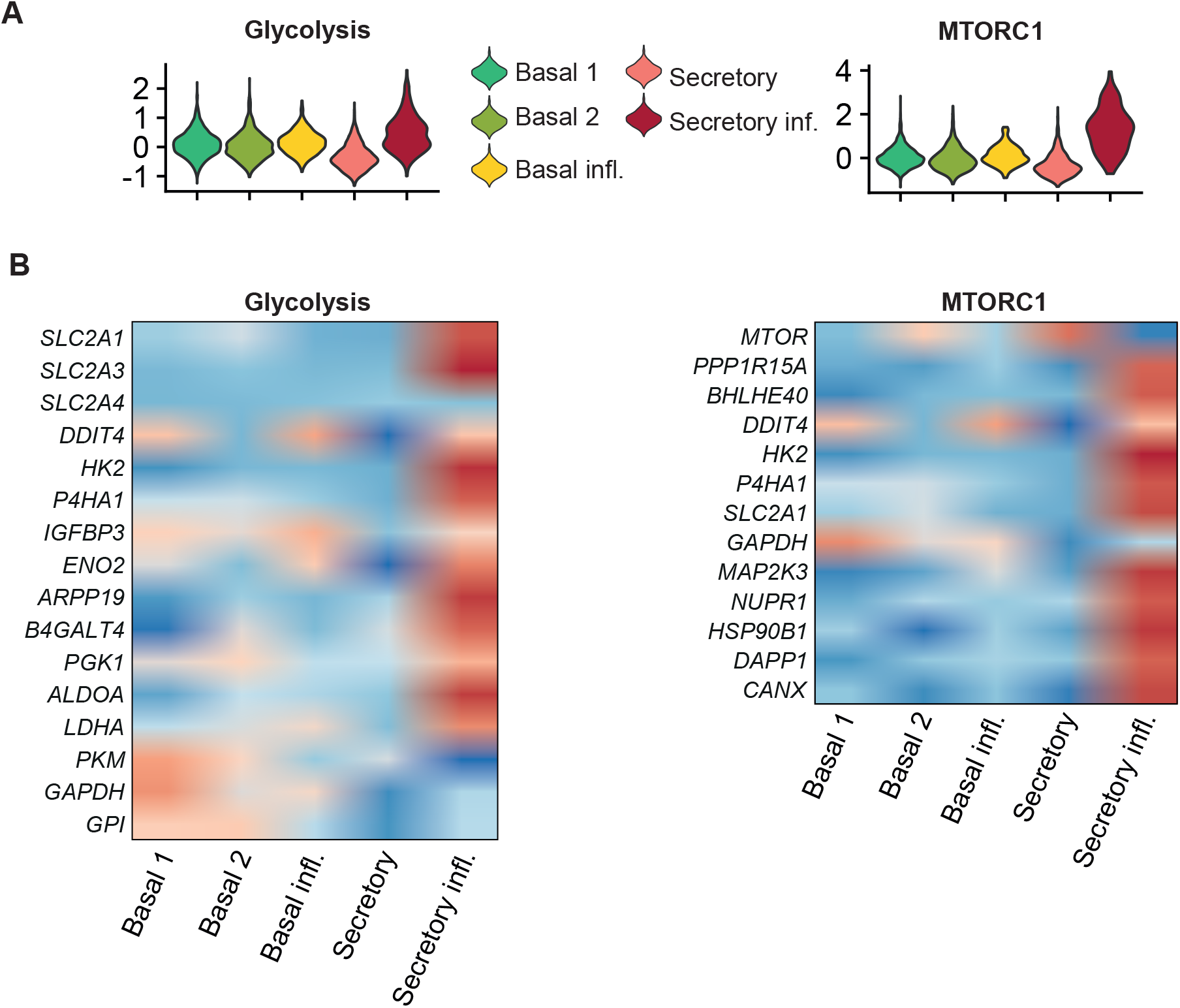
Secretory inflammatory cells have a distinct transcriptional profile of enhanced glycolysis and mTOR-signaling. **A**) Violin plots of the metabolic gene signature scores for glycolysis and the mTOR signaling pathway between the cell clusters. Other metabolic signature scores are depicted in supplemental figure S2. **B**) Heatmaps depicting the average scaled expression of genes related to glycolysis and the mTOR signaling pathway, split between the cell clusters.

### Rapamycin induces a specific proportional decrease of secretory inflammatory cells

We hypothesized that mTOR-mediated glycolysis was key for the expansion of secretory inflammatory cells during stimulation with flagellin. To investigate whether mTOR-inhibition would affect this flagellin-induced change, we next compared HBE cells stimulated with flagellin in the presence or absence of rapamycin. We applied the same approach as in Figure 1, and arrived at seven cell clusters which we annotated based on the expression of canonical genes (see **Supplemental Table 1**). Again, we identified several states of basal cells and secretory cells, as well as ionocytes and ciliated cells (**Figure 4A**). The secretory subclusters included two clusters that mainly differed based on the expression of genes from the SERPIN family (*SERPINB3, SERPINB1, SERPINB4* and *SERPINB7*). Cells clustered as secretory 1 and secretory inflammatory expressed higher levels of SERPIN-related genes which are known for being involved in inflammatory and immune cellular responses^14^. Cells clustered as secretory 2 expressed the classical club cell markers^15^ including genes related to mucous cell differentiation (*TFF3* and *AGR2*) and growth factors related to host defense (*LCN2* and *BPIFB1*). The secretory inflammatory cluster showed high expression of genes coding for inflammatory mediators (*IL36G* and *CXCL8*) (see **Figure 4B** for the top 10 DEGs of each cluster). The two identified basal states included the canonical markers described in **Figure 1C**, with increased expression of inflammatory genes in the basal inflammatory compartment (*S100A8* and *S100A9*) (**Figure 4B**).

**Figure 4.**
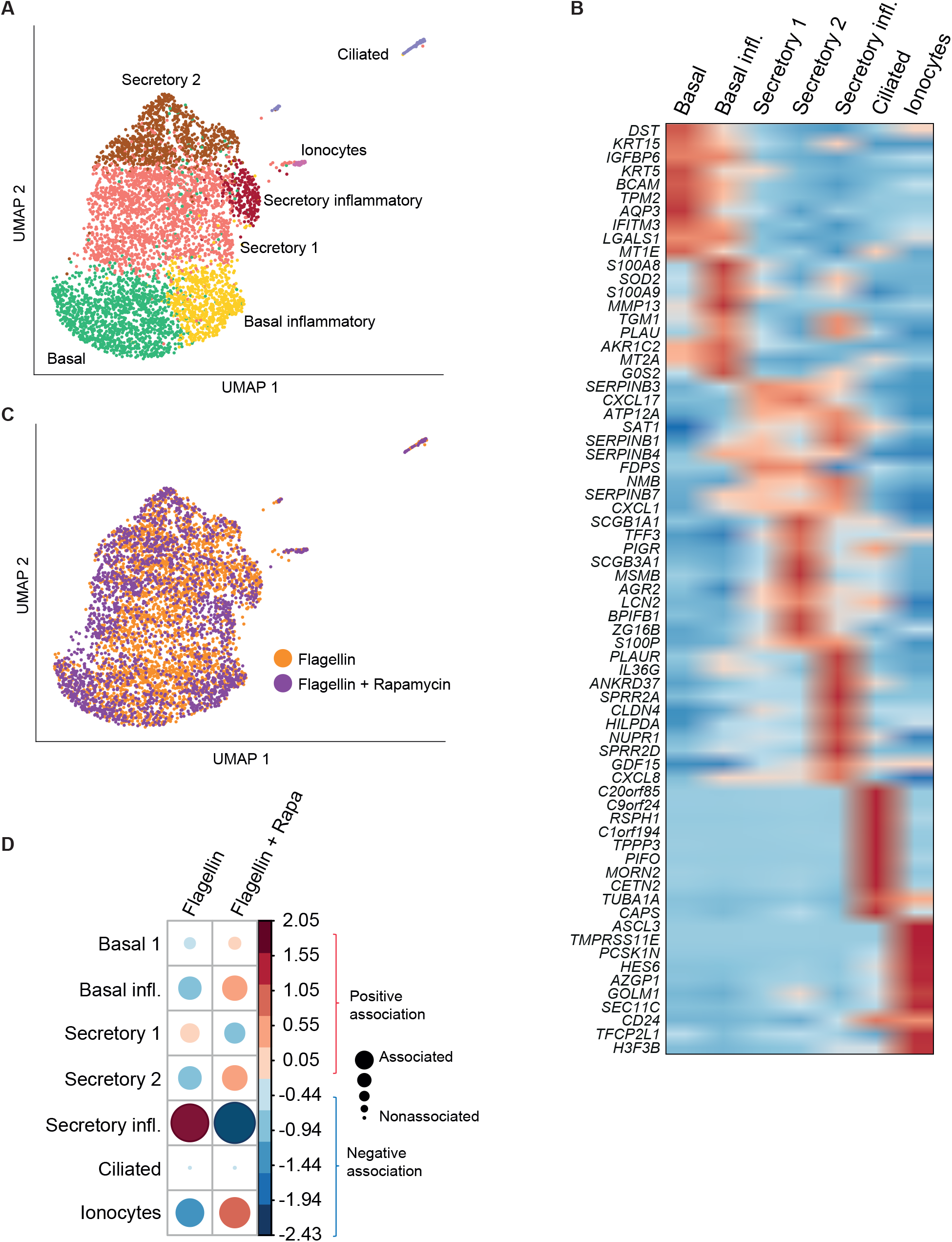
Rapamycin induces a specific proportional decrease of secretory inflammatory cells. **A**) UMAP of HBE cells, stimulated by either flagellin or flagellin + rapamycin, color indicates the cluster as identified by single-cell transcriptomic analysis. Each dot represents one cell. **B**) Heatmap showing the expression of the top differentially expressed genes (DEGs) for each cluster. Genes from the canonical markers used for cluster identification are listed in **Table S1. C**) Same as panel A, now colored by experimental condition. **D**) Correlation plot depicting cluster enrichment between the flagellin and flagellin + rapamycin condition. Dot size is proportional to the Pearson’s residual of the chi-squared test (reflecting the difference between the observed and expected proportion), while the color represents the degree of association from Pearson’s chi-squared residuals (red is a positive association, blue is a negative association).

Next, we assessed the impact of rapamycin on the flagellin-stimulated cells (see **Figure 4C** for the UMAP colored per condition). Strikingly, we observed a large proportional difference of the secretory inflammatory cluster: secretory inflammatory cells were clearly less abundant in the presence of rapamycin (**Figure 4D**). The other clusters showed only subtle differences – with slightly higher proportions of basal inflammatory cells, secretory 2 cells and ionocytes in the rapamycin condition – in line with our hypothesis that rapamycin may specifically limit expansion of secretory inflammatory cells. To analyze whether this specific proportional decrease of secretory inflammatory cells was indeed related to impaired mTOR-signaling and glycolysis, we next assessed the transcriptomic differences between the two conditions.

### Rapamycin induces a transcriptional profile of reduced glycolysis and inflammation, and decreases GLUT1 expression in secretory cells

Our group previously showed that rapamycin suppresses flagellin-induced glycolysis and mTOR signaling in bulk HBE cells^7^. Comparison of the metabolic and inflammatory gene signature scores induced by flagellin in the presence or absence of rapamycin within the secretory cell metacluster confirmed that rapamycin downregulated mTOR-signaling, glycolysis and inflammatory genes (**Figure 5A**), whilst other metabolic pathways were slightly upregulated (**Figure S3**). **Figure 5B** depicts key genes in glycolysis – akin to **Figure 3B** – for the distinct secretory cell clusters and split between the conditions, showing that rapamycin induced downregulation of glycolytic genes in every subset of secretory cells. *SLC2A1*, the gene encoding GLUT1, was among the top downregulated genes by rapamycin in flagellin-stimulated bulk HBE cells (data not shown) and in the glycolysis gene pathway in secretory cells (**Figure 5B**). Confocal microscopy confirmed enhanced expression of GLUT1 in secretory cells in HBE cultures stimulated with flagellin; inhibition of the mTOR pathway by rapamycin limited the abundance of GLUT1 in these cells, losing the organization observed in the flagellin-only condition (**Figure 5C**). Importantly, rapamycin treatment only resulted in a proportional decrease of secretory inflammatory cells, and not the other secretory subsets, as shown in **Figure 4D**. Together these data indicate that flagellin induces a proportional increase of inflammatory cell subsets in the human airway epithelium, of which specifically secretory inflammatory cells rely on mTOR-signaling and glycolysis. Incubation with rapamycin diminishes the upregulation of these transcriptomic pathways and results in a specifically decreased abundance of secretory inflammatory cells.

**Figure 5.**
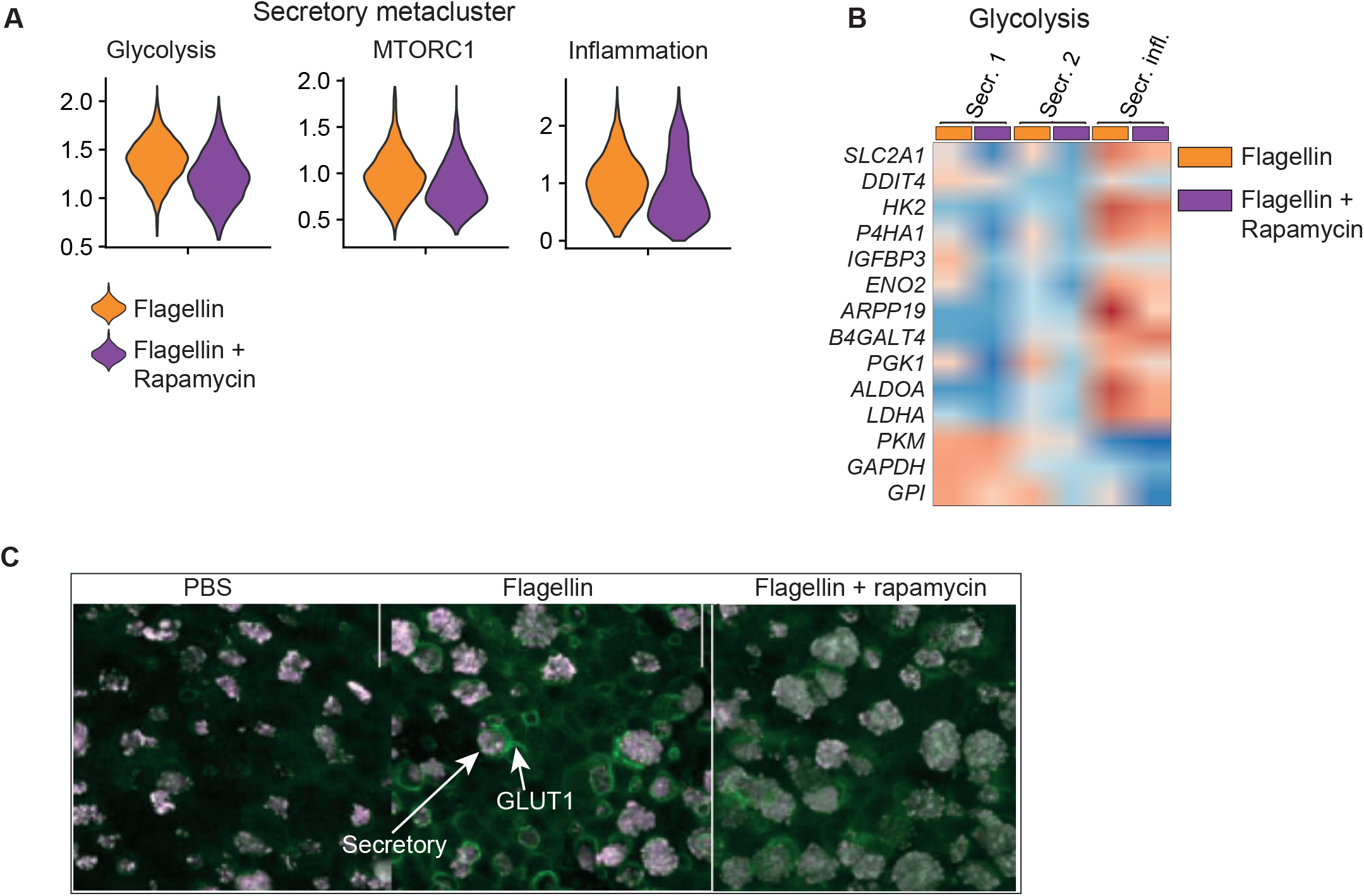
Rapamycin induces a transcriptional profile of reduced glycolysis and inflammation, and decreases GLUT1 expression on secretory cells. **A**) Violin plots of the gene signature scores for glycolysis, mTOR and inflammation between cells in the secretory metacluster, either stimulated with flagellin or flagellin + rapamycin. Scores for other metabolic signatures are shown in supplemental figure S3. **B**) Heatmap depicting the average scaled expression of genes related to glycolysis, split between the secretory cell clusters and the experimental condition (flagellin, or flagellin + rapamycin). **C**) Confocal microscopy showing HBE cells co-stained for GLUT1 (green) and secretory cells (purple; MUC5B^+^) in the indicated conditions.

## DISCUSSION

The respiratory epithelium plays a crucial role in innate defense against pathogens that invade the airways. We recently showed that the bacterial component flagellin activates HBE cells by increasing intracellular glycolysis through an mTOR dependent mechanism^7^. The airway wall consists of several cell types with distinct functions^2^ and we here argued that our primary HBE cell cultures growing at air-liquid interface may be a suitable model to study the involvement of different cell types in the metabolic rewiring and inflammatory responses induced by flagellin. We used scRNAseq to disclose seven cell clusters in HBE cells, of which flagellin specifically increased the abundance of inflammatory basal and inflammatory secretory cells. Remarkably, only inflammatory secretory cells displayed upregulation of genes involved in mTOR signaling and glycolysis in response to flagellin and inhibition of mTOR by rapamycin reversed this flagellin effect, as well as inflammatory gene transcription in this cell subset. Collectively, these data identify inflammatory secretory cells as the target of flagellin-induced glycolysis and inflammation in the airway epithelium.

Many studies that investigated the immune responses in respiratory epithelial cells made use of cell lines, which fail to recapitulate the cellular heterogeneity in the airway wall. This prompted us to use primary HBE cell cultures in air-liquid interface, which more resemble the cellular composition and physiological environment in the human airways, to study the metabolic regulation of mucosal immunology in the respiratory tract^7,8^. scRNAseq analyses have added an extra layer of information about the human airways, revealing distinct cell types that previously had not been recognized by conventional methods such as microscopy^2,12^. We here leveraged this technique to study cell-specific effects of flagellin in HBE cultures and the role of the mTOR pathway herein. Clear flagellin effects were detected in the inflammatory clusters within basal and secretory metaclusters, while ionocytes and ciliated cells were non-responsive. In accordance, the Human Protein Atlas (https://www.proteinatlas.org) reports the strongest expression of the gene encoding TLR5, the signaling receptor for flagellin, in basal and secretory cells in the human airways, with relatively low expression levels in ionocytes and ciliated cells. Our current data add to this that within the basal and secretory cell metaclusters, subclusters can be identified with distinct responsiveness to flagellin. Amongst these, inflammatory secretory cells responded to flagellin with an upregulation of genes regulating the glycolysis and mTOR pathways, and inhibition of the mTOR pathway by rapamycin strongly reduced the abundance of this cell subset and its inflammatory gene transcription, suggesting that our previous finding of flagellin-induced mTOR-driven glycolysis in HBE cultures is driven by inflammatory secretory cells^7^. Secretory (goblet) cells are important for key immune functions such as secretion of mucus and antimicrobial peptides^2^. As such, the current data shed light on the interaction between metabolism and mucosal immunity at single cell resolution.

scRNAseq analyses provide extraordinary “snapshot” information about the dynamic cellular composition of the airways. Distinct cell types identified by scRNAseq can represent cells at different differentiation stages. Pseudotime analyses, a computational method that orders cells according to their trajectory or lineage in time, have indicated that secretory cells can function as precursor for ciliated cells^16^. On the other hand, basal cells are considered multipotent stem cells from which other major subpopulations, including secretory and ciliated cells, originate^17^. Hence, our current results suggest that cells contained within the human airways can acquire or lose metabolic programs that support their inflammatory properties during their trajectories of differentiation.

Knowledge of the interaction between cellular metabolism and the capacity to mount an energy demanding inflammatory response is mainly derived from studies in lymphocytes and myeloid cells^3–7^. In these cells mTOR activation can have distinct effects, which may relate to differences in experimental conditions and the intrinsic complexity of the mTOR pathway. The present study, taken together with our previous report using bulk HBE cells^7^, points at a proinflammatory immune stimulating role of mTOR in flagellin-stimulated HBE cells. In agreement, we showed that flagellin administered via the airways of mice increased the expression of epithelial-cell specific immune mediators such as *Cxcl1, Cxcl2* and *Csf3* in bronchial brushes by an mTOR dependent mechanism^7^.

Flagellin is part of the bacterial flagellum, a surface filament important for bacterial motility^9^. Flagellated bacteria are sensed in the airway lumen at the apical site of the mucosa through TLR5, which initiates a brisk immune response. The potency of flagellin to elicit an innate response in the respiratory epithelium has made it an attractive candidate for an immune enhancing strategy in the airways during pneumonia^9^. The present study shows that flagellin specifically targets cellular subsets within the human airways, and that flagellin-induced proinflammatory effects in inflammatory secretory cells are regulated by the mTOR pathway. These findings advance our understanding of the regulation of mucosal immunology during infections with flagellated bacteria and provide novel information on how flagellin modulates immune functions in the airways.

## METHODS

### HBE cells and stimulation

HBE cells were derived from healthy tracheobronchial tissue obtained from a lobectomy in a patient with lung cancer at the Amsterdam University Medical Centers in the Netherlands, as previously reported^7,8^. The Institutional Review Board of Amsterdam UMC approved the study protocol (2015-122#A2301550) and written informed consent was obtained from the donor before sampling. HBE cells were isolated according to Fulcher’s protocol^18^. In short, passage 2 (P2) to P4 passaged cells were differentiated in 24-well Transwell inserts (Corning, Corning, NY, USA) coated with human type IV placental collagen in submerged PneumaCult-Ex Plus media (StemCell Technologies, Vancouver, Canada). When cells reached confluency, the media was replaced by PneumaCult-ALI medium (StemCell Technologies) on the basolateral side and the apical side was exposed to the air, forming an air-liquid interface. The basolateral media were renewed every two or three days for around 30 days. For cell stimulation experiments, flagellin from *Pseudomonas aeruginosa* (1 μg/ml; Invivogen, Toulouse, France) was added to the apical compartment and left for 24 hours. For mTOR inhibition experiments, rapamycin (10 nM; Cayman Chemical, Ann Arbor, MI, USA) was added to the medium 1 hour prior to flagellin stimulation. After stimulation, HBE cells were digested for 30 min at 37°C in the presence of 5 mg/ml Liberase TM (Sigma; St. Louis, MO, USA) and 10 mg/ml DNase (Roche; Basel, Switzerland). Fixable Viability Dye kit (eBioscience; San Diego, CA, USA) was used to assess cell viability by FACS analysis (> 90% in all samples).

### Single-cell library generation and sequencing

The Chromium Single Cell 5’ Library & Gel Bead Kit v1.1 (10X Genomics, Pleasanton, CA) was used to generate the libraries following the manufacturer’s instructions. In short, we sorted 200,000 single cells per condition. Each sample was labelled with TotalSeq c human hashtag antibodies (Supplemental Table 2). Samples were incubated for 30 minutes with 1 μl of antibody per sample. Thereafter, the antibody was washed three times with PBS 0.1% BSA. Cells were counted and 50,000 cells from each sample were pulled together. The pooled samples were loaded in the 10X chromium and loaded the cells on the 10X machine. Libraries were generated according to standard protocol (Chromium Next GEM Single Cell V(D)J Reagent Kits v1.1 Rev E; 10X Genomics). Libraries were sequenced using the Illumina HiSeq4000 (Illumina, San Diego, CA, USA). Each position from the 10X chip was loaded into 1 HiSeq 4000 lane.

### Data analysis

All libraries were aligned using CellRanger 3.1 (10x Genomics). Cell deconvolution and analysis were performed using the R package ‘Seurat’^19^. The cells were filtered based on number of features (nfeature_RNA), number of genes (nCount_RNA), and percentage of mitochondrial reads (percent.mt) using Seurat ‘subset(object, subset=nCount_RNA>1000 and nCount_RNA<10,000 and nFeature_RNA>200 and percent.mt<20)’. We calculated cell cycle scores as follows: (‘CellCycleScoring(Healthy_object_Cycle), s.features=s.genes, g2m.features=g2m.genes, set.ident=TRUE’) using a list of S phase and G2M phase genes preloaded in Seurat. The cells were then scaled and normalized using the function ‘SCTransform(object, vars.to.regress=c(‘nCount_RNA’, ‘percent.mt’, ‘S.Score’, ‘G2M.Score’)). Principal components for each set of cells as shown in individual figures were identified using the ‘RunPCA’ function. Cells were then further processed for clustering and visualization using the Seurat functions ‘FindNeighbors’, ‘FindClusters’, and ‘RunUMAP’. The number of principal components used as input was determined by using the ‘ElbowPlot’ function and identifying the number of the PCs which explain most of the data variance. Differential expression analysis was performed using the functions ‘FindAllMarkers’ or ‘FindMarkers’ and the following parameters: ‘min.pct=0.25, logfc.threshold=0.25, assay=‘SCT’’.

### Statistical analysis

All analyses were performed in R version 4.1.2. Enrichment analysis was performed by first generating contingency tables with the cell distributions per cluster. Next, chi-squares for the contingency tables were calculated and correlation plots were generated with the R package *corrplot*. Comparisons with at least more than two residuals of difference were considered biologically relevant. All gene expression analyses were corrected for multiple testing using the Benjamini-Hochberg method, with significance defined as an adjusted p<0.05. Pathway analysis was performed using Gene Set Enrichment Analysis based on the Molecular Signatures Database (MSigDB)^13^.

### Confocal microscopy

Cells were fixed in 4% formaldehyde for 20 min and stored submerged in PBS at 4°C until analysis. HBE were permeabilized for 1 hour in 0.1% Triton-X followed by blocking in 1% BSA in PBS for 30 min. Samples were stained with the following primary antibodies: MUC5B (sc-20119; Santa Cruz Biotechnology; Dallas, TX, USA), p63 (AB124762; Abcam, Cambridge, UK) p63 (AF1916; R&D Systems; Minneapolis, MN, USA), ß-tubulin-Cy3 (C4585; Sigma) and Glut1-A488 (ab195359; Abcam). The following secondary antibodies were used: anti-rabbit-AF488 (A21202; LifeTechnologies; Carlsbad, CA, USA), anti-rabbit-Cy5 (711/175-152; Jackson Immuno Research Labs; West Grove, PA, USA) and anti-goat-AF488 (Ab150157; Abcam). The nuclei were stained using the ProLong™ kit (ThermoFisher Scientific; Waltham, MA, USA). Cell imaging was performed on a Leica SP8 confocal microscope (Leica; Wetzlar, Germany). Images are shown as maxi-projection on the z-stack planes.

## Data availability

The accession number for the single-cell RNA sequencing data reported in this paper is GEO: GSE (*not yet available*).

## ACKNOWLEDGEMENTS

Ivan Ramirez-Moral was funded by the Era-Net JPIAMR/ZonMW (grant 50-52900-98-201). Part of this work was funded by European Union H2020 (FAIR project, grant 847786). The authors thank Marja E. Jakobs (Core Facility Genomics, Amsterdam-UMC) for her technical support during the single cell RNA sequencing pipeline and Daisy Picavet and Ron Hoebe (Cellular Imaging Core Facility, Amsterdam-UMC) for helping in the acquisition and analysis of confocal microscopy images.

## AUTHOR CONTRIBUTIONS

Conceptualization: I.R-M. and T.v.d.P. Methodology: I.R-M., K.d.H, S.v.L., X.Y. and F.A.V.B. Validation: I.R-M., A.R.S., J.M.B. and F.A.V.B. Formal analysis: I.R-M., A.R.S., J.M.B., and F.A.V.B. Investigation: I.R-M., and F.A.V.B. Resources: T.v.d.P. and M.D.d.J. Writing - Original draft: I.R-M. A.R.S. and T.v.d.P. Writing - review and editing: I.R-M., A.R.S., J.M.B., F.A.V.B., A.F.d.V., K.d.H., S.v.L., X.Y., M.D.d.J. and T.v.d.P. Visualization: I.R-M., A.R.S., J.M.B. and F.A.V.B. Funding acquisition: T.v.d.P.

## Supplementary Material

**Supplemental Table 1.** Human airway epithelial cell canonical markers used for scRNA-seq clustering.

| Cell type | Selected canonical genes | Proposed function |
| --- | --- | --- |
| Basal 1 cells | KRT5<sup>high</sup>, KRT15 | Mature stem cells involved in regeneration and repair of the epithelium |
| Basal 2 cells | KRT5<sup>low</sup> | Suprabasal cells intermediate between basal and club cells |
| Secretory cells (goblet cells) | SCGB1A1, MUC5AC, MUC5B, SCGB3A1 | Secretion of mucus, antimicrobial peptides and immune-mediators |
| Ciliated cells | FOXJ1, PIFO | Clearance of mucus |
| Ionocytes | FOXI1 | Ion transport, fluid and regulation of pH |

**Supplemental Table 2.**
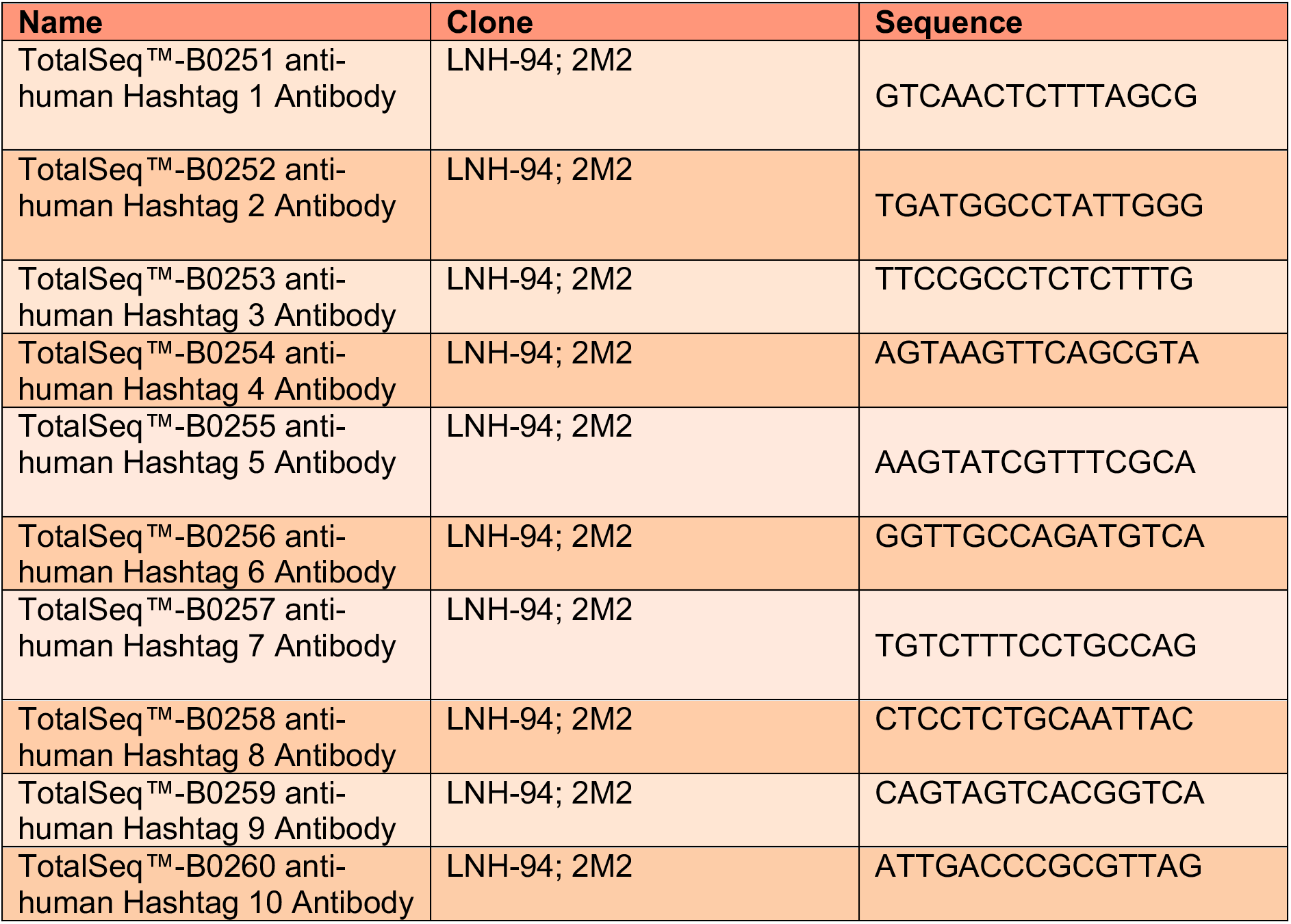
Hashtag antibodies sequences.

**Figure S1.**
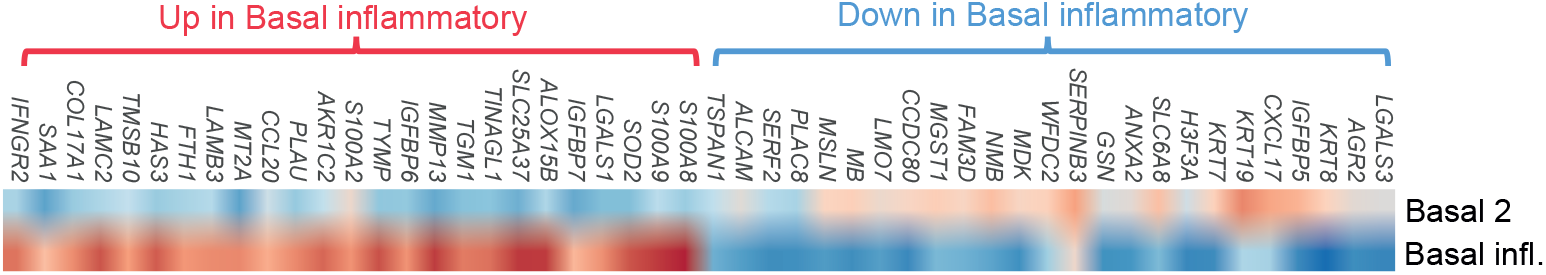
Transcriptional differences between the cell subsets. Heatmap showing the expression of the top 25 increased and top 25 decreased DEGs derived from comparing the basal inflammatory and basal 2 cells (log fold change > 0.25, adjusted p<0.05).

**Figure S2.**
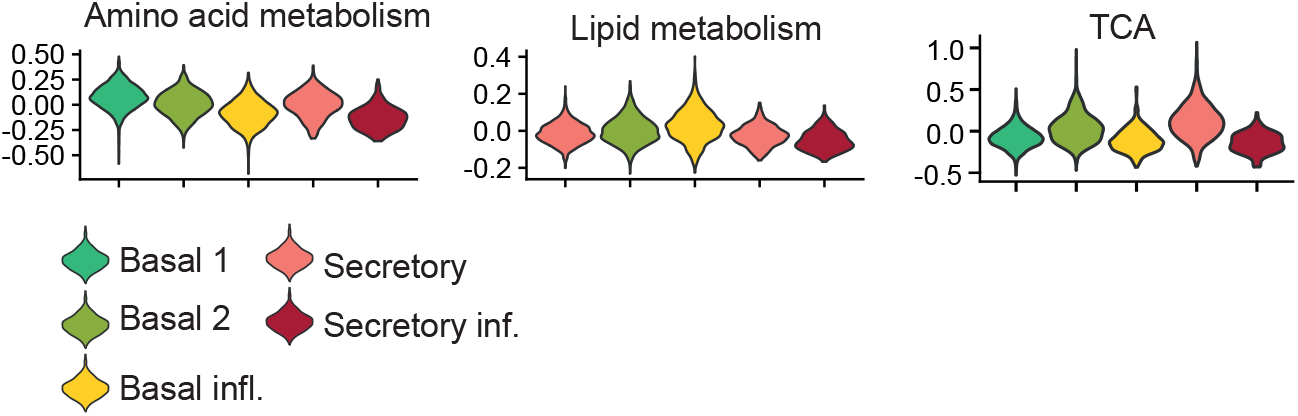
Metabolic signature scores between cell subsets. Violin plots of metabolic gene signature scores between the cell clusters.

**Figure S3.**
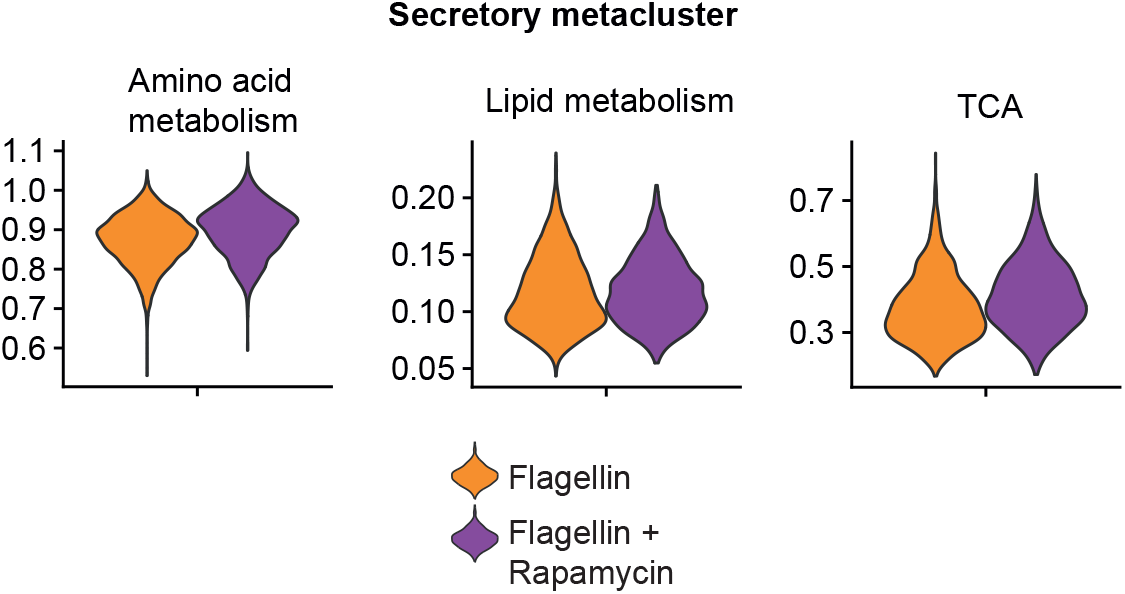
Metabolic signature scores in secretory cells. Violin plots of metabolic gene signature scores for all secretory cells split between the conditions: stimulated with flagellin or stimulated with flagellin and rapamycin.

